# The probability distribution of the reconstructed phylogenetic tree with occurrence data

**DOI:** 10.1101/679365

**Authors:** Ankit Gupta, Marc Manceau, Timothy Vaughan, Mustafa Khammash, Tanja Stadler

## Abstract

We consider a homogeneous birth-death process with incomplete sampling. Three successive sampling schemes are considered. First, individuals can be sampled through time and included in the tree. Second, they can be occurrences which are sampled through time and not included in the tree. Third, individuals reaching present day can be sampled and included in the tree. Upon sampling, individuals are removed (i.e. die).

The outcome of the process is thus composed of the reconstructed evolutionary tree spanning all individuals sampled and included in the tree, and a timeline of occurrence events which are not placed along the tree. We derive a formula allowing one to compute the joint probability density of these, which can readily be used to perform maximum likelihood or Bayesian estimation of the parameters of the model.

In the context of epidemiology, our probability density allows us to estimate transmission rates through a joint analysis of epidemiological case count data and phylogenetic trees reconstructed from pathogen sequences. Within macroevolution, our equations are the basis for taking into account fossil occurrences from paleontological databases together with extant species phylogenies for estimating speciation and extinction rates. Thus, we provide the theoretical framework for bridging not only the gap between phylogenetics and epidemiology, but also the gap between phylogenetics and paleontology.

## 1. Introduction

Birth-death processes are used extensively in both epidemiology and macroevolution, to model the underlying population dynamics of, respectively, infected individuals and species. For simplicity, we will refer to the atomic particles of the process as *individuals* in this paper. In its most simple form, a birth-death process describes the population dynamics of a set of independent individuals, each of which can give birth to another individual with a constant birth rate *λ*, or die with a constant death rate *µ*.

Seminal results on this process have been derived by Kendall (1948), who already evoked potential applications to epidemiology. Much more recently, Nee et al. (1994) and Nee and May (1997) have made important developments to the theory, showing how to compute the probability density of the *reconstructed evolutionary tree*, i.e. the tree obtained by first tracing all genealogical relationships between individuals, before erasing all branches that do not reach present. This has paved the way to extensive use of birth-death processes in modern phylogenetics, with many refinements: birth and death rates have been proposed to vary through time (Morlon et al., 2011; Stadler, 2011), to vary across lineages in the tree (Alfaro et al., 2009), to vary depending on the *type* of individuals (Maddison et al., 2007), or to depend on the number of individuals (Etienne et al., 2012; Leventhal et al., 2013).

In order to fit various applications, the sampling scheme of individuals has also been put under careful scrutiny. Nee et al. (1994) initially suggested that individuals could be sampled at present with a given probability *ρ*; this uniform sampling is also called *field of bullets* sampling. Yet, trees that were reconstructed using this sampling scheme were always ultrametric trees describing the genealogical relationships between present-day individuals only. This assumption was relaxed by Stadler (2010), who additionally modeled the sampling of individuals throughout the process, with a fixed per-individual sampling rate *ψ*. When sampled, an individual is displayed along the tree and the reconstructed (non-necessarily ultrametric) tree corresponds to the genealogical relationships between all sampled-through-time and sampled-at-present individuals. This model opened the way to numerous new applications in epidemiology and macroevolution, allowing to take into account non-synchronous data. In phylogenetics, this process is commonly used as a prior on the genealogical relationships between individuals sampled through time (Stadler et al., 2011). In macroevolution, it allows one to use molecular and paleontological evidence together, by simultaneously considering present-day species and the subset of fossil taxa which can be placed along a tree using morphological characters (Zhang et al., 2015; Gavryushkina et al., 2017).

One step further towards considering even more data jointly in one analysis has been performed by Vaughan et al. (2019), who introduce two types of sampling through time. The first one, which is named *sampling and sequencing*, is the same as previously described. The term *sequencing* here referring to the fact that the individual is placed unambiguously along the reconstructed tree using its genetic sequence. Alternatively, an individual could be *sampled and not sequenced*, in which case its existence is only recorded as a time point occurrence along a timeline. This approach is very promising, for it enables one to consider jointly data from case count epidemiological studies together with trees reconstructed from pathogens sequenced during an outbreak. Alternatively, in the context of macroevolution, it allows one to use poorly preserved fossil occurrences, which could not be placed along the reconstructed tree. Yet, in its current form, the inference framework proposed by Vaughan et al. (2019) relies on computer intensive Monte-Carlo simulations within a particle filtering approach which prevents its use on large datasets. Similarly, (Heath et al., 2014) proposes placing occurrences on a fixed tree using Markov chain Monte Carlo methodology. Here the drawback is again the computer intensive approach as well as relying on a fixed tree rather than sequences.

In this study, we derive a closed form formula for the joint probability density of a reconstructed tree with individuals sampled through time and a record of occurrences, i.e. sampling times for individuals not included in the tree. The underlying model is a birth-death process with sampling through time and at present, where upon sampling individuals are removed (i.e. die). The density can readily be included within phylodynamic tools as a prior in a Bayesian inference framework based on sequences and occurrences, or can be used for maximum likelihood parameter estimation based on a tree and occurrences. Its computational efficiency opens the way to analyse large datasets available for either epidemiology or macroevolution studies.

The aim of the paper is to provide a mathematical study of the birth-death model giving rise to serially sampled reconstructed trees and occurrences. We first introduce model notations and some preliminary observations. Then, we provide an alternative derivation of the probability density of the reconstructed evolutionary tree under the birth-death process with sampling through time (Stadler, 2010), assuming that an individual is removed upon sampling. Finally, we show how to extend this derivation to account for non-sequenced occurrences, and provide an analytical formula to compute the probability density of the reconstructed tree and occurrences. We finally discuss the use of this density to perform inferences in epidemiology and macroevolution in a maximum likelihood or Bayesian setting.

## 2. Model and notations

We consider a constant rate birth-death process with incomplete sampling. We allow the possibility that for some sampled individuals, the attachment times within the tree are known, whereas for others only their sampling times is known but their attachment times within the tree are unknown. The latter sampling events are called *occurrences*.

We assume that an individual gives birth at rate *λ*, and its death rate is (*µ* + *ψ* + *ω*). Here *µ* is the death rate *without sampling, ψ* is the death rate *with sampling and tree-inclusion* and *ω* is the death rate *with sampling but without tree-inclusion* (i.e. an occurrence). Additionally we assume that extant individuals, alive at present time, are sampled and included in the reconstructed tree with probability *ρ*. For convenience, we will refer to the five types of events in this model using their parameter names. In other words, *λ*-events refer to the creation of a new individual, *µ*-events refer to death events without sampling, *ψ*-events refer to death events with sampling and tree inclusion, *ω*-events refer to death events with sampling but without tree inclusion, and finally *ρ*-events refer to the sampling of extant lineages at present time.

Time is assumed to be 0 at present, and increases going into the past. The process starts at time *t*_*or*_ > 0 in the past, with one infected lineage and it generates a birth-death lineage tree from which the reconstructed tree is derived. Our data consists of both the reconstructed tree 𝒯 and the occurrence set 𝒪. The reconstructed tree 𝒯 is generated by all the *ψ*- and *ρ*-events giving rise to leaves, together with all *λ*-events subtending those leaves. The occurrence set 𝒪 consists of the times of all the *ω*-events, all belonging to the interval (0, *t*_*or*_); see Figure 1 for an example.

**Figure 1:**
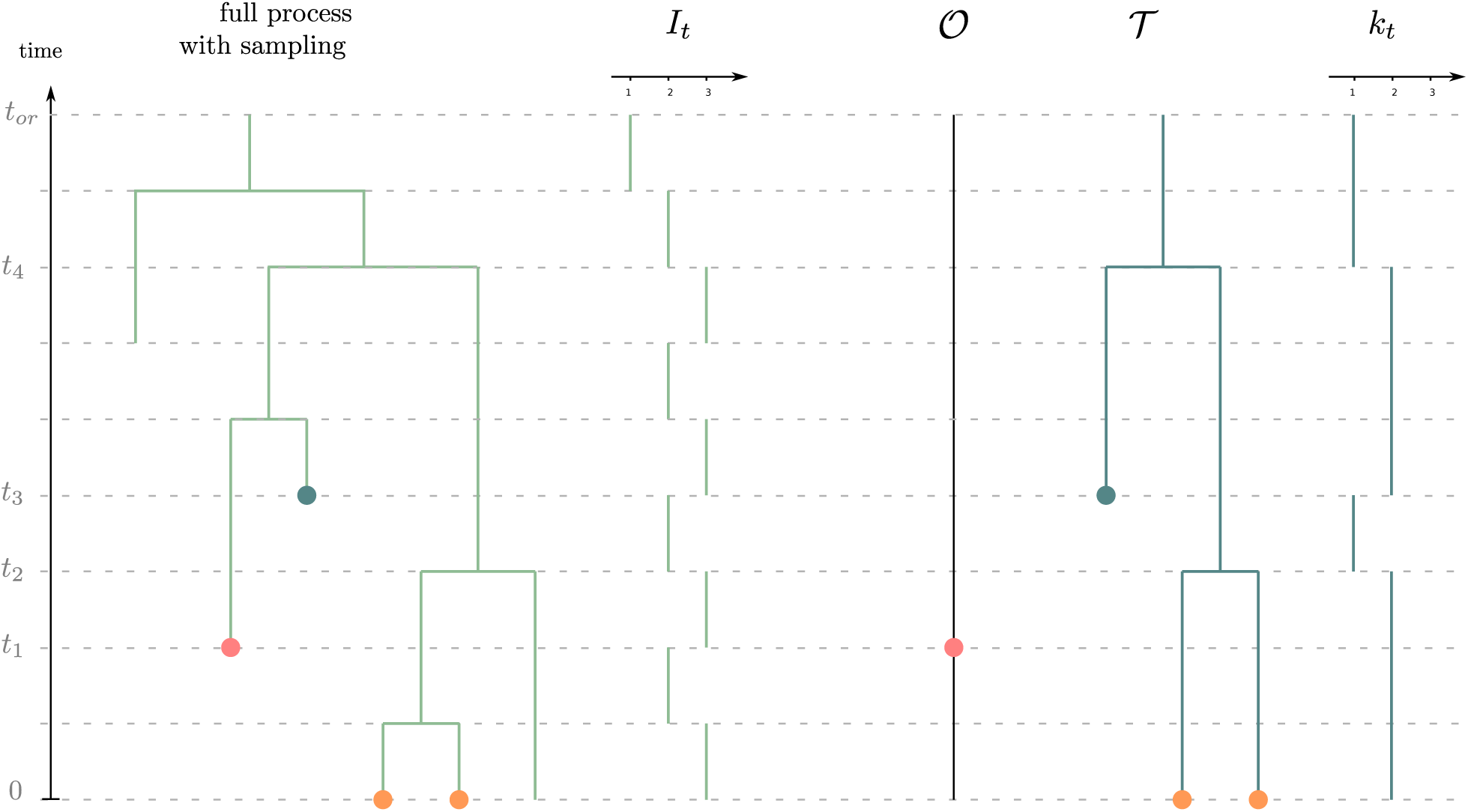
General setting of the method. On the left in green, the full process with sampling is shown. Red dots correspond to *ω*-sampling (sampling through time without sequencing), blue dots correspond to *ψ*-sampling (sampling through time with sequencing) and orange dots correspond to *ρ*-sampling at present. On the right, the observations are shown: a sampled-through-time reconstructed tree 𝒯 together with sequential observations 𝒪 of sampled individuals along a timeline.

We will be interested in the probability density of the joint observation of (𝒪, 𝒯), which will ultimately depend on the times at which observed *λ*-, *ψ*-, and *ω*-events happen, but not on the tree topology (see e.g. (Aldous et al., 2001; Stadler, 2010)). All these times pooled together are denoted as *t*_0_, *t*_1_, *…, t*_*n*_, starting with *t*_0_ = 0 and ending with *t*_*n*_ = *t*_*or*_, and the number of observed lineages on the interval (*t*_*h*_, *t*_*h*+1_) is denoted *k*_*h*_.

### 2.1. Introducing useful quantities

Let *u*(*t*) be the probability that an individual alive at time *t* before today has no sampled extinct or extant descendant lineages. Also let *p*(*t*) be the probability that a lineage alive at time *t* before today has precisely one sampled extant lineage and no sampled extinct descendant lineages. We can write the Master Equations for these two probabilities as

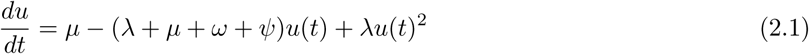

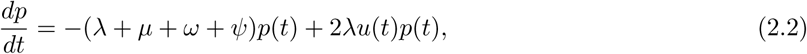

with initial condition (*u*(0), *p*(0)) = (1 − *ρ, ρ*). Defining

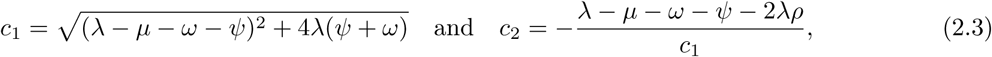

the solution of these Master Equations (see Theorem 3.1 in Stadler (2010)) is given by

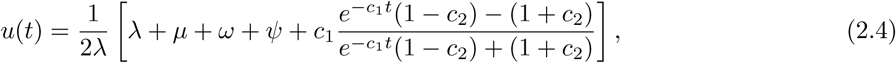

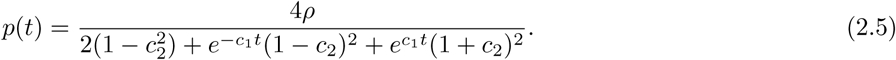

### 2.2. Key first observations

In order to understand better future calculations, we must examine the birth-death dynamics of the *number of hidden lineages* in greater detail. Suppose (*X*_*s*_)_*s*≥0_ be the *forward in time* stochastic process describing the number of hidden lineages in some time-interval [*s*_1_, *s*_2_] ⊂ [0, *t*_*or*_], where *s*_1_ *< s*_2_ and we use this new letter *s* because time is oriented (only in this section of the manuscript) from the origin 0 towards the present *t*_*or*_. The number of observed lineages in 𝒯 is fixed and equal to *k* in this interval and also there is no occurrence event in this interval. This stochastic process (*X*_*s*_)_*s*≥0_ is a birth-death process over nonnegative integers ℕ_0_ and it has the following characteristics:

- When the state is *i*, the rate of birth is *λ*(*k* + *i*) and if this event happens, the state moves to (*i* + 1) with probability

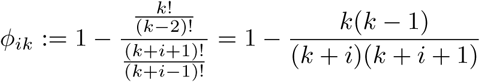

and with probability 1 − *φ*_*ik*_ the state moves to some *absorbing state* Δ outside the state space ℕ_0_ = {0, 1, *…*}. Note that as the number of infected lineages increases from *k* + *i* to (*k* + *i* + 1) upon a birth event *φ*_*ik*_ is simply the probability that when we “look backwards”, the two coalescing lineages are not both among the sampled lineages. In case the two coalescing lineages are among the sampled lineages then the number of observed lineages will not be fixed in the time-interval [*s*_1_, *s*_2_] and hence this birth-death trajectory for total number of infected lineages becomes infeasible, which is equivalent to saying that it gets absorbed at state Δ.
- When the number of hidden lineages is *i*, the rate of death is (*µ* + *ψ* + *ω*)(*k* + *i*) and if this event happens for some state *i* > 0 then the state moves to (*i* − 1) with probability

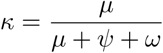

and it moves to the absorbing state Δ with probability (1 − *κ*). Note that (1 − *κ*) is simply the probability of a death event either being a *ω*-event or a *ψ*-event. Both such events will violate our assumption that there is no occurrence event and the number of observed lineages is fixed in the time-interval [*s*_1_, *s*_2_]. Moreover if *i* = 0 and a death event happens then the birth-death trajectory again becomes infeasible and so it gets absorbed at state Δ.

It is clear that due to the presence of an absorbing state outside the state space, the process (*X*_*s*_)_*s*≥0_ is *non-conservative*. The rate of change of the distribution of this process can be specified by its generator (see Chapter 3 in Ethier and Kurtz (1986)). We can define this operator on test functions *f*: ℝ_+_ × ℕ_0_ → ℝ by

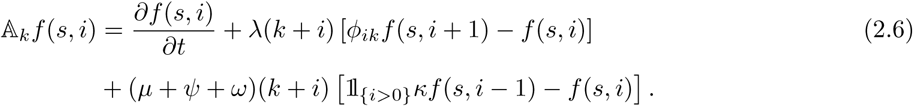

Here we implicitly assume that function *f* is continuously differentiable in the first coordinate. We now come to a very important proposition on which our whole analysis depends.

**Proposition 2.1.** *Let u*(*t*) *be given by* (2.4) *and define the function f*_*k*_: ℝ_+_ × ℕ_0_ → ℝ *by*

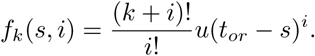

*Then the action of generator* 𝔸_*k*_ *on function f*_*k*_ *simplifies to*

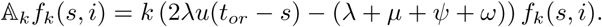

*Moreover the following is a positive martingale in the interval* [*s*_1_, *s*_2_] *w.r.t. the filtration generated by process X,*

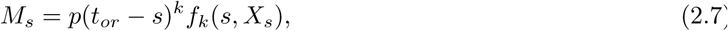

**Remark 2.2.** *The function f*_*k*_ *is a time-varying eigenfunction for the operator* 𝔸_*k*_ *and the corresponding eigenvalue is k* (2*λu*(*t*_*or*_ − *s*) − (*λ* + *µ* + *ψ* + *ω*)).

**Proof** See Appendix. □

Note that the fact that *M*_*s*_ is a martingale implies that

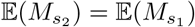

which yields

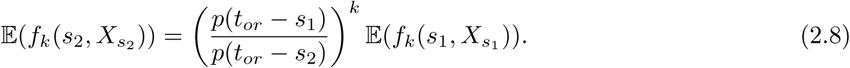

Reversing the direction of time and letting *t*_1_ = (*t*_*or*_ − *s*_2_) and *t*_2_ = (*t*_*or*_ − *s*_1_), conditioning on 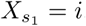, and exploiting the time-homogeneity of the Markov process (*X*_*s*_)_*s*≥0_ we can express (2.8) as

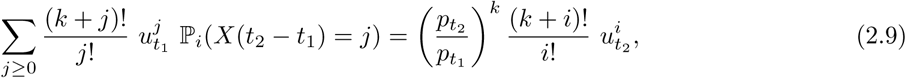

where the subscript *i* denotes that the initial state is *X*_0_ = *i*. This formula will help us provide an alternative derivation of the probability density of a reconstructed tree without occurrence data originally introduced by Stadler (2010).

## 3. Revisiting the probability density of the reconstructed tree with samples through time

We assume in this section that *ω* = 0, and there is no occurrence data. We wish to offer an alternative derivation of the probability density of the reconstructed tree spanning both *ρ*-sampled and *ψ*-sampled individuals as in Stadler (2010).

Let *t*_0_ *< t*_1_ *< t*_2_ *< … < t*_*n*_ be the ordered set of times at which the tree events occur backward in time starting from the present time *t*_0_ = 0 and culminating at *t*_*n*_ = *t*_*or*_. For any *t* ∈ [0, *t*_*or*_], call *k*_*t*_ the number of observed lineages at time *t* in 𝒯 and let 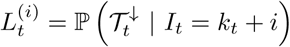 be the probability for the observed tree *below* time *t* (i.e. in the time-interval [0, *t*]) when the total number of lineages is *k*_*t*_ + *i* at time *t*. Clearly the probability density of the observed tree is 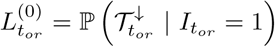, and to obtain this quantity we would like to study how *L*_*t*_ evolves with time *t* and how it depends on the state *i*.

We now introduce an *ansatz* for the form of 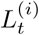, given by

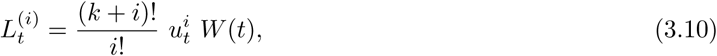

where *W* (*t*) is some real-valued *weight* function that only depends on time *t* but not on *i* or *k*.

Let us consider the interval (*t*_*h*−1_, *t*_*h*_) for some *h* ≥ 1. In this interval the number of observed lineages is constant and equal to *k*. Using ansatz (3.10) and applying the Markov property we obtain

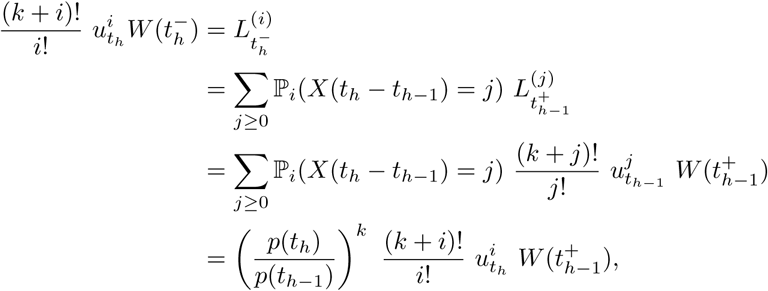

where the last equality follows from (2.9). This proves that under ansatz (3.10)

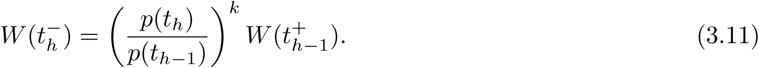

We now examine how *W* (*t*^+^) and *W* (*t*^−^) are related, for any time *t* at which we observe an event. Suppose first that the event at time *t* is a *ψ*-event. Then *k*^+^ = *k*^−^ + 1 and if the total number of lineages is *k*^+^ + *i* at time 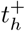 then the rate at which this event happens is *ψ*(*k*^+^ + *i*) and subsequently the total number of lineages falls to (*k*^−^ + *i*). This shows that

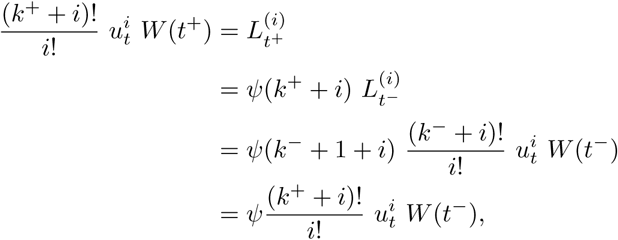

and hence

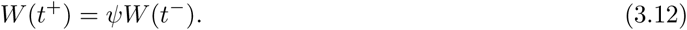

Now suppose the event at time *t* is a *λ*-event. Then *k*^+^ = *k*^−^ − 1 and if the total number of lineages is (*k*^+^ + *i*) at time *t*^+^ then the rate at which this event happens is *λ*(*k*^+^ + *i*) and with probability 1*/*((*k*^+^ + *i*)(*k*^+^ + 1 + *i*)) this event is a coalescent event between two observed ordered lineages. Once this event happens the total number of lineages increments to (*k*^−^ + *i*). Therefore

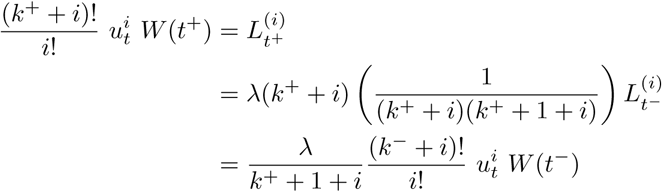

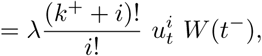

which proves that

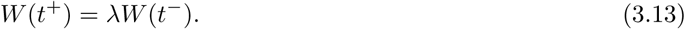

Let *k*_0_ be the number of extant lineages at time *t*_0_ = 0. Using relations (3.11), (3.12) and (3.13) we can propagate the weight function *W* (*t*) backward in time starting from 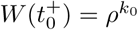 and ending at 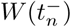 which is equal to the tree density ℒ (𝒯). This backward propagation scheme is described in Algorithm 1 and it yields a closed-form formula very similar to theorem 3.5 in Stadler (2010). Since individuals are here removed upon sampling, our result can be derived from theorem 3.5 by setting *k* = 0 and dropping *p*_0_(*y*_*i*_) factors corresponding to the death of *ψ*-sampled leaves.

### Algorithm 1 Computes the probability density ℒ (𝒯)

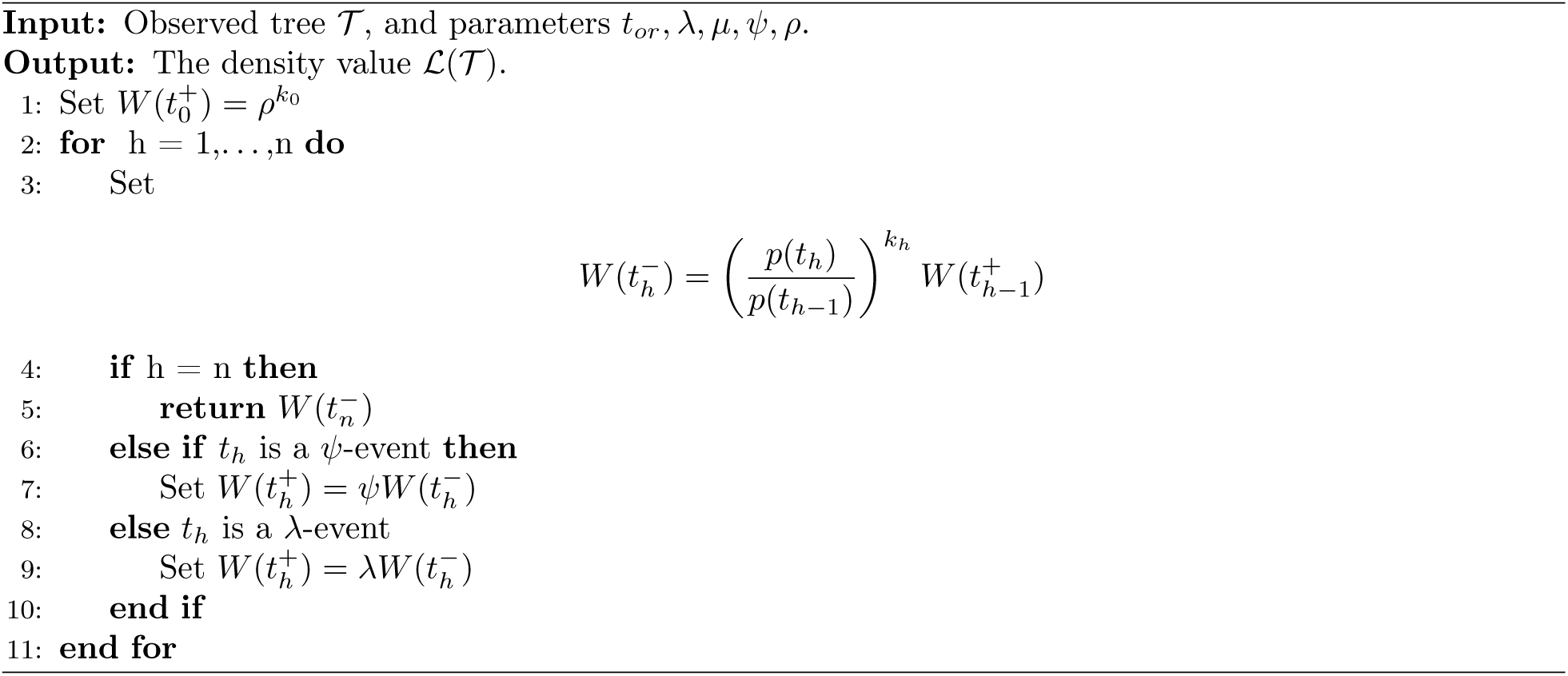

## 4. The density of the reconstructed tree and case count record

We now consider the scenario where *ω* ≠ 0 and we have occurrence data *O* along with the observed lineage tree 𝒯. As in the previous section, let *t*_0_ *< t*_1_ *< t*_2_ *< … < t*_*n*_ be the ordered set of times at which the events occur backward in time starting from the present time *t*_0_ = 0 and culminating at *t*_*n*_ = *t*_*or*_. For any *t* ∈ [0, *t*_*or*_], let 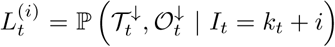 be the probability for the observed lineage tree and occurrence data *below* time *t* (i.e. in the time-interval [0, *t*]) when the number of observed lineages in 𝒯 is *k*_*t*_ at time *t* and the number of hidden lineages is *i*. Clearly the probability density of the observed tree and occurrence data is again 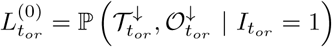, and to obtain this quantity we would like to study how *L*_*t*_ evolves with time *t* and how it depends on the state *i*. It will become evident that ansatz (3.10) will not work for this probability, and so we generalize this ansatz as

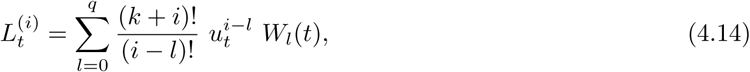

where *q* is a nonnegative integer, *k* is the number of observed lineages at time *t*, and *W* (*t*) = (*W*_0_(*t*), *…, W*_*q*_(*t*)) is some vector-valued *weight* function that only depends on time *t* but not on *i* or *k*. Note that for *q* = 0, this ansatz becomes the previous ansatz (3.10).

### 4.1. Backward evolution at punctual events

We first examine how *W* (*t*^+^) and *W* (*t*^−^) are related when *t* is an event time. Suppose first that the event at time *t* is a *ψ*-event. Then *k*^+^ = *k*^−^ + 1 and if the total number of lineages is *k*^+^ + *i* at time *t*^+^ then the rate at which this event happens is *ψ*(*k*^+^ + *i*) and subsequently the total number of lineages falls to (*k*^−^ + *i*). This shows that

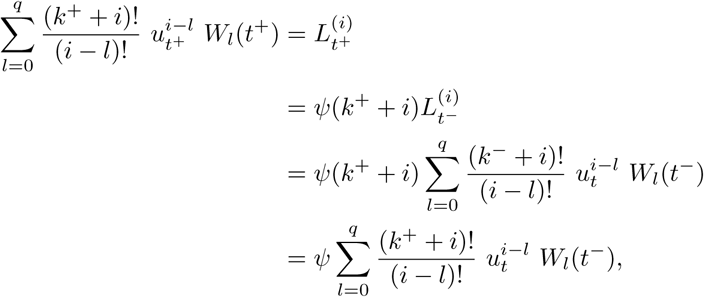

and hence

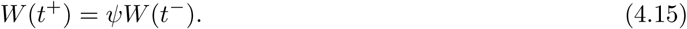

Now suppose that the event at time *t* is a *λ*-event. Then *k*^+^ = *k*^−^ − 1 and if the total number of lineages is (*k*^+^ + *i*) at time *t*^+^ then the rate at which this event happens is *λ*(*k*^+^ + *i*) and with probability 1*/*((*k*^+^ + *i*)(*k*^+^ + *i* + 1)) this event is a coalescent event between two observed ordered lineages. Once this event happens the total number of lineages increments to (*k*^−^ + *i*). Therefore

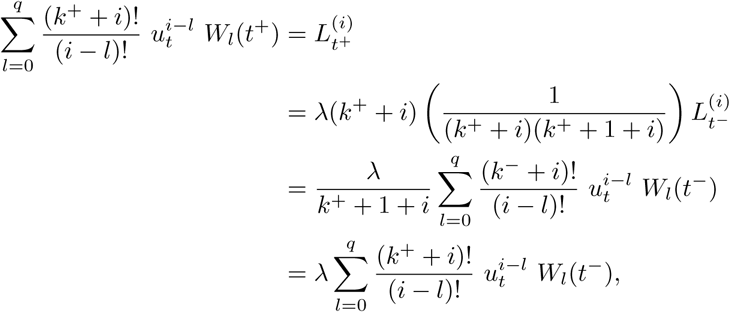

which proves that

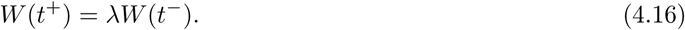

So far the transition conditions (4.15) and (4.16) are identical to what we encountered in the previous section. However the difference comes for *ω*-events as we now discuss. Suppose *t* is a *ω*-event. Then *k*^+^ = *k*^−^ and if the total number of lineages is (*k*^+^ +*i*) at time *t*^+^ then the rate at which this event happens is *ω*(*k*^+^ +*i*) and subsequently the total number of lineages falls to (*k*^+^ + *i* − 1). This shows that

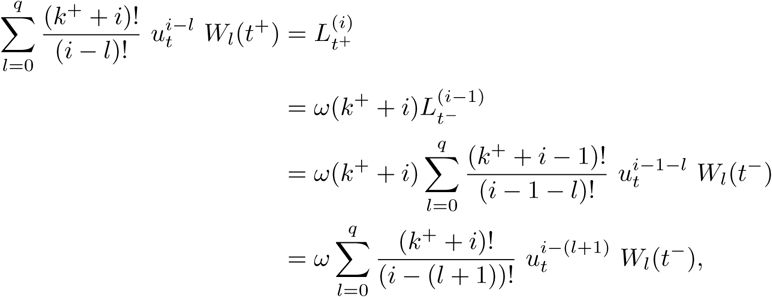

and hence

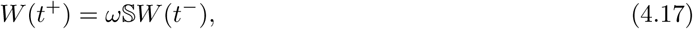

where 𝕊 is the shift-operator defined by

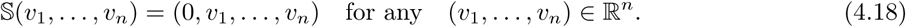

Setting *q* = 0 in this calculation shows that the simpler ansatz (3.10), in which *W* (*t*) is a scalar function instead of a vector-valued function, is not compatible with the requirement that *W* (*t*) is not a function of *i* or *k*.

### 4.2. Backward evolution on a time interval without punctual events

Now that we have the transition conditions, (4.15), (4.16) and (4.15), we can propagate the weight function *W* (*t*) backward in time, provided we can evaluate how it evolves in a time-interval (*t*_*h*−1_, *t*_*h*_) for any *h* ≥ 1, and this backward evolution preserves our ansatz (4.14). In order to study this backward evolution we need some new notation and a simple lemma, which we now provide.

For any time *t* ≥ 0 and *θ* ∈ (0, 1), define

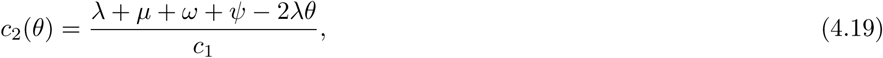

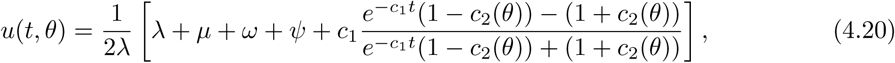

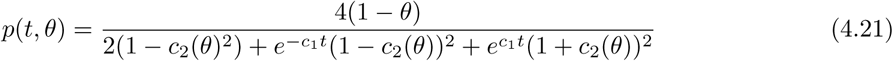

and where *c*_1_ is as given in (2.3). Notice that if we set *θ* = 1 − *ρ* then *c*_2_(*θ*), *u*(*t, θ*) and *p*(*t, θ*) become identical to *c*_2_, *u*(*t*) and *p*(*t*) defined in Section 2. Henceforth for any *k* ∈ ℕ_0_ we also define the ratio

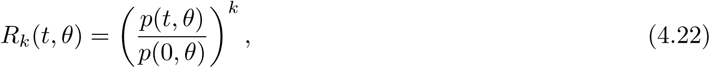

and when *k* = 1, we drop the subscript and refer to *R*_*k*_(*t, θ*) as *R*(*t, θ*). To ease further algebra, we also name here the function corresponding to the denominator of *p*(*t, θ*), namely *q*(*t, θ*) = 4(1 − *θ*)*/p*(*t, θ*). The next lemma gives us analytical expressions for the higher order partial derivatives of *R*_*k*_(*t, θ*) and *u*(*t, θ*) w.r.t. *θ*.

**Lemma 4.1.** *Let u*(*t, θ*) *and R*_*k*_(*t, θ*) *be as defined by* (4.20) *and* (4.22) *respectively. Then for any n* = 1, 2, *… we have the following:*

*(A) Let* 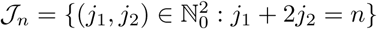. *Then* 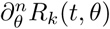 *is given by*

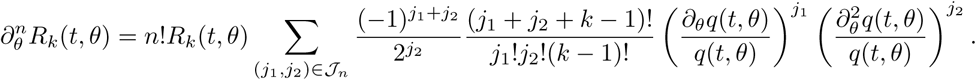

*(B) The quantity* 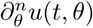 *is given by*

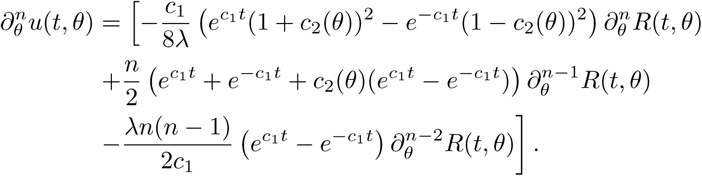

**Proof** See Appendix. □

Recall the formula (2.9), for the probability evolution on a time-interval [*t*_1_, *t*_2_] on which the number of observed lineages remains fixed at *k*, and call now *θ* = *u*(*t*_1_, 1 − *ρ*). Appealing to the semi-group property of solutions to ODEs (2.1)-(2.2) we get

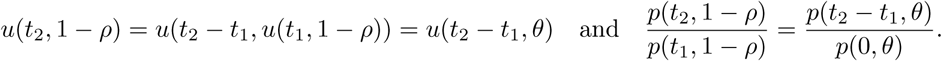

Hence we can rewrite (2.9) as

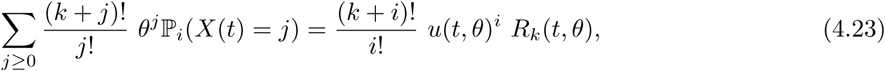

where *t* = (*t*_2_ − *t*_1_). Note that the probability ℙ_*i*_(*X*(*t*) = *j*) does not depend on *θ*. Differentiating (4.23) *l* times w.r.t. *θ* we obtain

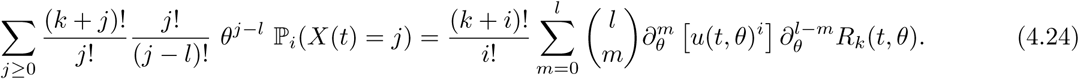

Applying the *Faà di Bruno’s* formula (see Fraenkel (1978)), for any *m* = 1, 2, *…* yields

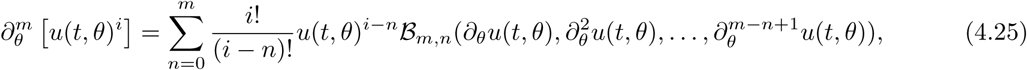

where ℬ*m, n*(*x*_1_, *x*_2_, *…, x*_*m*−*n*+1_) is the incomplete Bell polynomial. Such polynomials can be computed efficiently via a recurrence relation

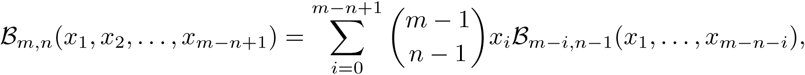

where ℬ_0,0_ = 1 and ℬ_*m,*0_ = ℬ_0,*m*_ = 0 for each *m* ≥ 1. Henceforth we denote

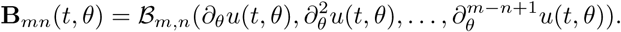

Substituting (4.25) in (4.24) we obtain

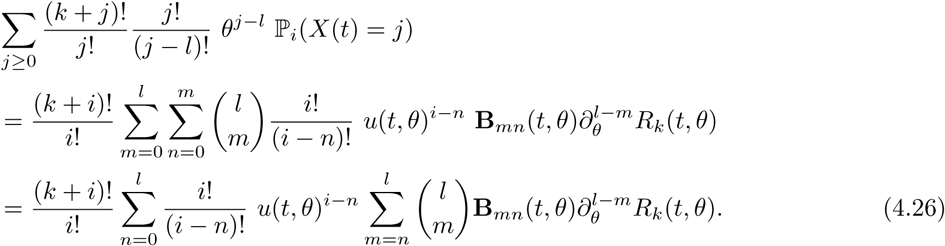

Letting

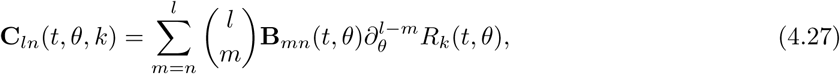

we can express (4.26) as

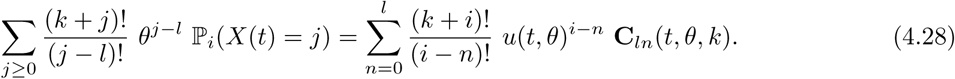

This formula will play a critical role in determining the backward propagation of the vector-valued weight function *W* (*t*) = (*W*_0_(*t*), *…, W*_*q*_(*t*)) between the transition points.

Indeed, let us consider the interval (*t*_*h*−1_, *t*_*h*_) for some *h* ≥ 1. In this interval the number of observed lineages is constant and we assume that it is equal to *k*_*h*_. Also let *θ*_*h*_ = *u*(*t*_*h*_, 1 − *ρ*) for each *h*. Using ansatz (4.14) and applying the Markov property we obtain

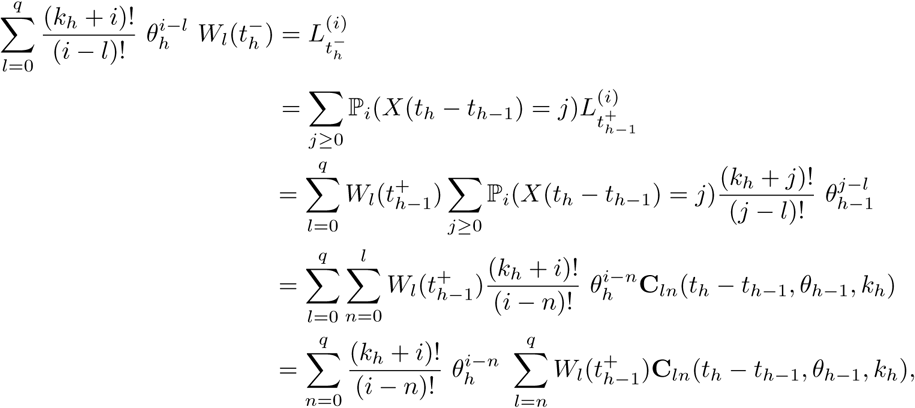

where the second-last equality is due to formula (4.28). This calculation proves that ansatz (4.14) is preserved by the backward evolution of the weight vector *W* (*t*) and

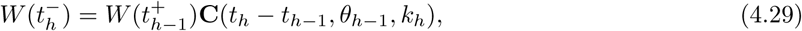

where 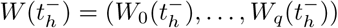 and 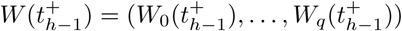 are (*q* + 1)-dimensional row vectors and **C**(*t*_*h*_ − *t*_*h*−1_, *θ*_*h*−1_, *y*_*h*_) is the (*q* + 1) × (*q* + 1) lower-triangular matrix matrix whose entries are given by **C**_*ln*_(*t*_*h*_ − *t*_*h*−1_, *θ*_*h*−1_, *k*_*h*_) for *l* ≥ *n* and 0 for *l < n*. Note that by using Lemma 4.1, this matrix can be analytically computed.

### 4.3. Summary of the likelihood computation

Let *k*_0_ be the number of extant and sampled lineages at time *t*_0_ = 0. Using relations (4.29), (4.15), (4.16) and (4.17) we can propagate the vector-valued weight function *W* (*t*) backward in time starting from 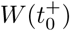 whose entries are all equal to 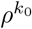 and ending at 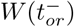, whose first component 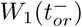 is equal to the target probability density ℒ(𝒯, 𝒪). Note that due to the presence of the shift operator S in (4.17) the dimensional of *W* (*t*) is (*q* + 1) where *q* is the number of occurrence events in the time-interval [0, *t*]. This backward propagation scheme is described in Algorithm 2 and it simplifies to Algorithm 1 in the absence of occurrence data.

#### Algorithm 2 Computes the probability density ℒ(𝒪, 𝒯)

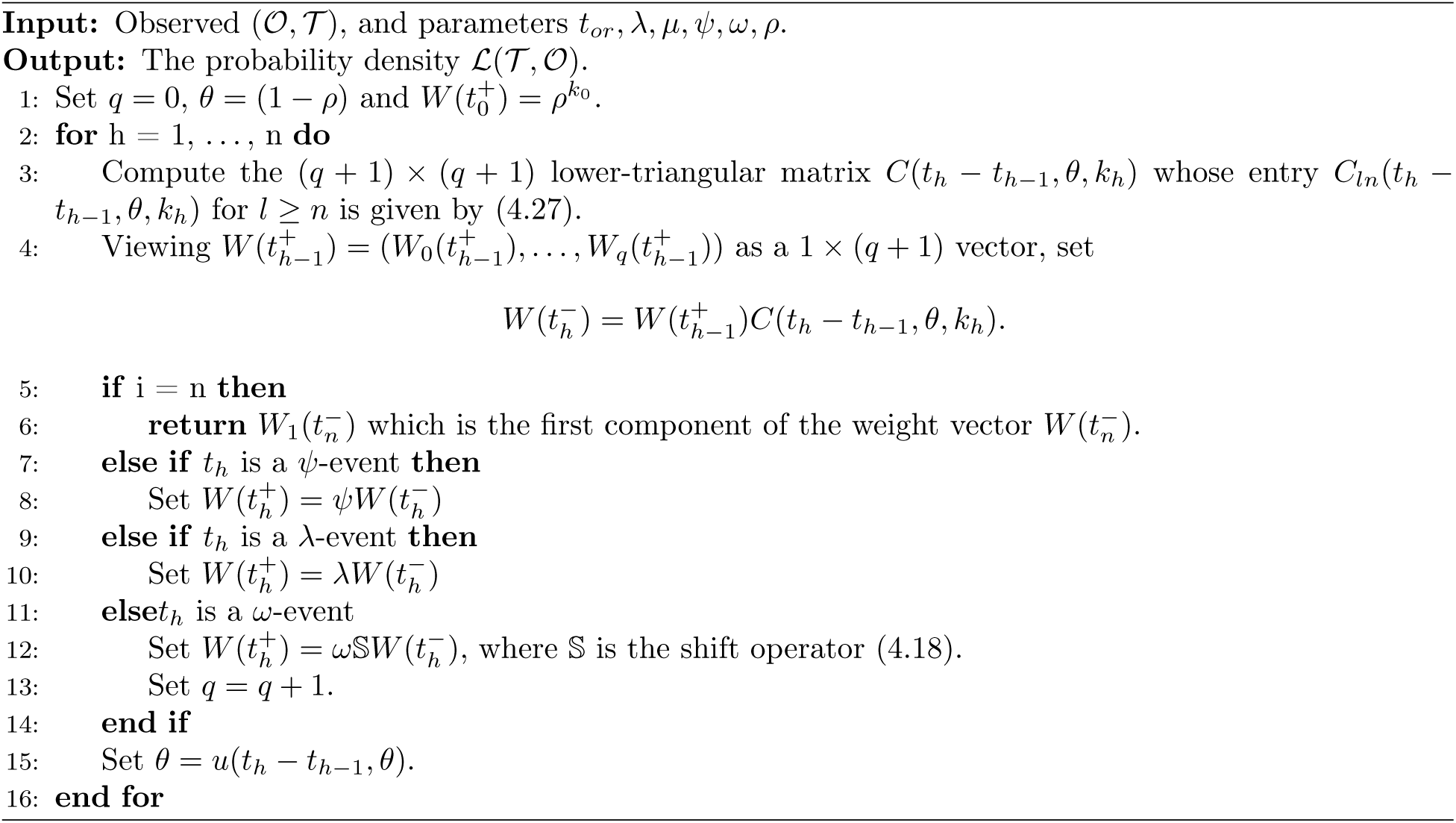

### 4.4. Numerical implementation and sanity check

The main algorithm to compute the probability density of (𝒪, 𝒯) has been implemented numerically and is available on GitHub: https://github.com/ankitgupta83/tree-and-occurrences/. As a check that the method leads to correct values, we compared the results obtained with our analytical calculation (Algorithm 2) with the ones obtained using Monte Carlo simulations as described in Vaughan et al. (2019) and implemented here in C++.

We consider two toy examples depicted in Figure 2(A) for which we calculate the probability density. The comparisons are shown in Figure 2(B) and Figure 2(C), and one can see that there is a close match between numerical values computed analytically and those estimated with simulations.

**Figure 2:**
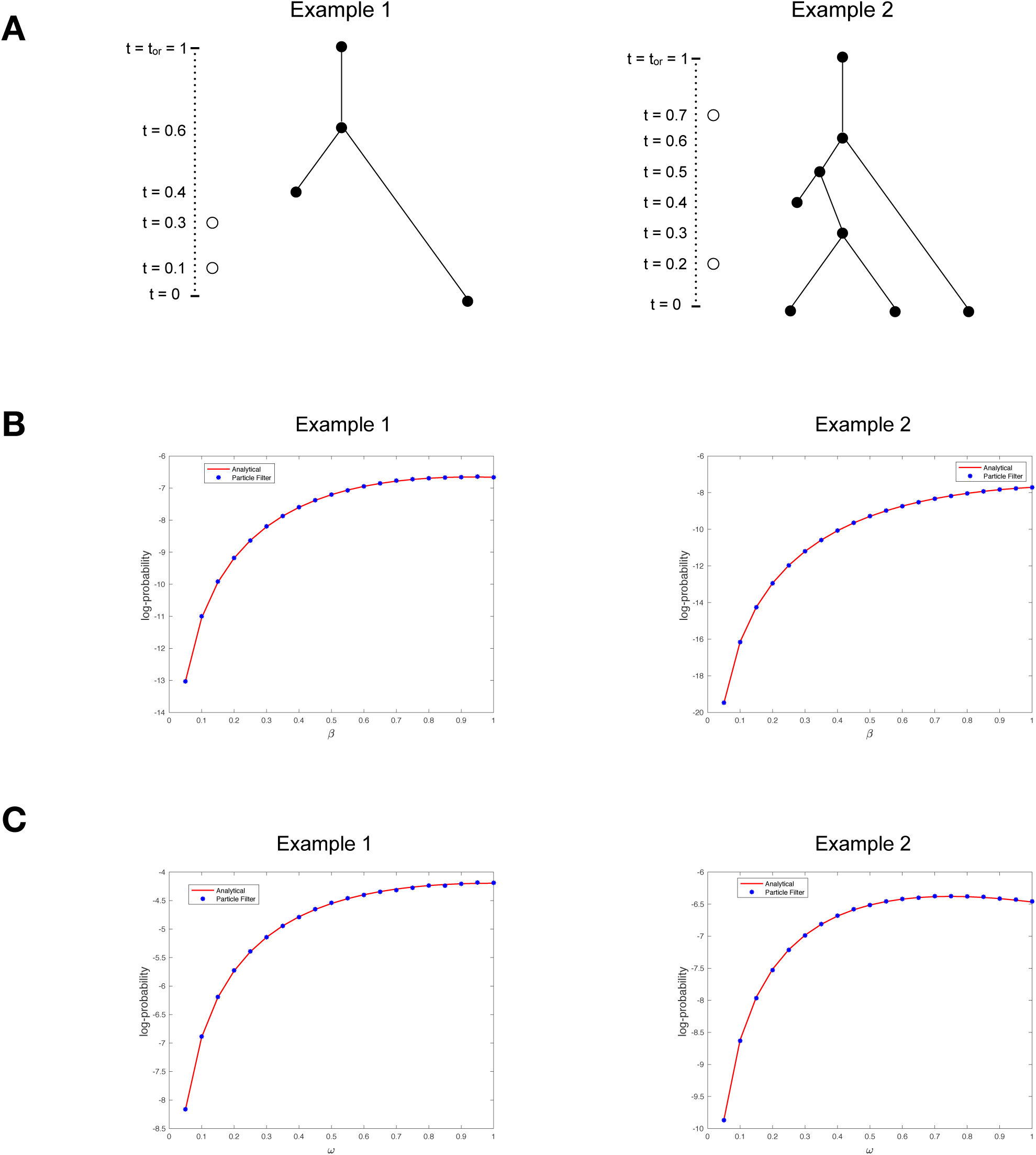
(A) Two toy examples of reconstructed trees along with occurrence data (open circles). (B) Comparison between probability densities obtained analytically with Algorithm 2 and the corresponding probability densities obtained with simulations as described in Vaughan et al. (2019). The parameter *λ* varies between 0 and 2, while the other parameters are set to *ψ* = 0.1, *µ* = 0, *ω* = 0.1 and *ρ* = 0.7. (C) Same as (B) except now the parameter *ω* varies between 0 and 1, while the other parameters are set to *ψ* = 0.1, *µ* = 0, *λ* = 1.25 and *ρ* = 0.7.

## 5. Applications

Our results pave the way to different applications that we present in this section.

### 5.1. Different flavors of the likelihood

Following Stadler (2010), we first point out that our main result, Algorithm 2, allows us to compute the density of the observations, given a model with fixed time of origin *t*_*or*_. This quantity, that we call *L* below, can be seen as an extension of Theorem 3.5 in Stadler (2010) (while assuming that samples correspond to death events), when occurrences can be observed.

Depending on the analysis that one wants to perform, it might be desirable to condition this density on various events, including,

1. the number of *ρ*-sampled leaves at present,

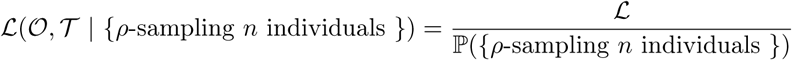

where the denominator is a well-known quantity which can, e.g., be found in Theorem 3.3 in Stadler (2010), replacing *µ* by *µ* + *ψ* + *ω* in our case.
2. the survival of the process up to the present,

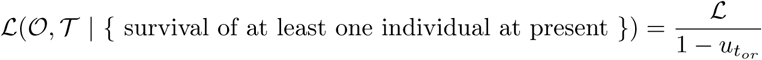
3. the survival of two lineages starting at time *t*_*mrca*_,

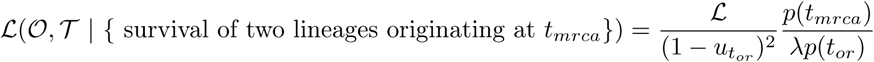

In each case, since the event we condition on is in the observed data, one only needs to divide by the probability of the event. These are only three possibilities that one could use. In the context of phylodynamics, conditioning on the survival of the process, or on the survival of two lineages starting at *t*_*mrca*_ is common practice, since we study an evolutionary process precisely because it survived.

We also stress that the probability density provided in this manuscript applies to an oriented and unlabelled tree. If one wishes to compare the probability densities of trees under different generating models, then it might be useful to add a combinatorial factor for dealing, e.g. with unoriented and labelled trees, as described in Stadler (2010).

### 5.2. Maximum likelihood estimators

The previous probability densities can readily be used to estimate parameters of the model from reconstructed trees and occurrences. Taken as a function of parameters, the density is indeed called likelihood, and *maximum likelihood estimators* can be obtained by maximizing the likelihood function over the parameters. Since it does not appear straight-forward to identify optimal parameter configurations analytically, we suggest relying on numerical optimizers instead. In this context, computing quickly the likelihood value using Algorithm 2 in place of the more computationally intensive Monte-Carlo algorithm proposed by Vaughan et al. (2019), can prove essential, since the optimization requires many calls to the function.

### 5.3. Bayesian analysis

This density is also a key component when using this model in a Bayesian framework, where one can estimate the parameters of the model directly from the occurrences and the sequencing data rather than fixing a t ree. Suppose we observe the sampling times of occurrences 𝒪 (individuals without any measurement data), the sampling times and genotypic or phenotypic measurements of *m ψ*-sampled individuals, and the genotypic or phenotypic measurements of *n ρ*-sampled individuals. We call 𝒜 the sequencing data, summarizing all the genotypic or phenotypic measurements (e.g. a nucleotide alignment of pathogens, or a collection of morphological traits for fossil species, or any combination of those), and *θ* all parameters relating to the model of character evolution along a tree.

Loosely defining *f* to represent all probability densities involved, and relying on the name of the random variable to know which one we refer to, one is generally interested in sampling from,

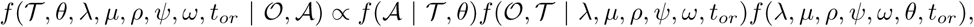

with *f* (*λ, µ, ρ, ψ, ω, θ, t*_*or*_) being the prior distribution on the model parameters. Standard MCMC (Markov-Chain Monte-Carlo) algorithms can be used to sample from this posterior distribution. As a result, from sequencing and occurrence data, we can directly obtain the marginal distribution of birth-death parameters integrated over the posterior distribution of trees.

### 5.4. Simulations of the process

While the raw process is quite easy to simulate forward-in-time, it can be more difficult to simulate it under a different conditioning event *C*, relating for example to the number of *ρ*-, *ψ*-, or *ω*-samples. Naive rejection sampling is not an efficient option if one wishes to condition on an event happening with very low probability. Adapting the particle filtering algorithm developed by Vaughan et a l. (2019) would b e a much better option.

Another option is to use the results of this paper and directly sample from the density *f* (𝒪, 𝒯 | *λ, µ, ρ, ψ, ω, t*_*or*_, 𝒞), using our ability to evaluate *f* (𝒪, 𝒯 | *λ, µ, ρ, ψ, ω, t*_*or*_) quickly, within an MCMC algorithm with any reasonable movement proposal changing the reconstructed tree and possibly occurrence times or numbers while satisfying the constraint 𝒞.

## 6. Discussion

In this study, we derived a closed-form probability density formula of a reconstructed tree and an occurrence record (i.e. a record of when cases occurred) under a linear birth-death model with sampling. This can readily be used for statistical purposes, to infer the parameters of the model in a maximum likelihood or a Bayesian framework.

In the context of epidemiology, this study offers a way to improve the accuracy of statistical estimates of key epidemiological parameters, such as the transmission and recovery rate (*λ* and (*µ* + *ψ* + *ω*)), and thus the basic reproductive number *R* = *λ/*(*µ* + *ψ* + *ω*), using jointly the occurrence record and the phylogenetic tree reconstructed from pathogen sequences. This should be of use to health policy makers as well as epidemiologists, enabling them to jointly use the epidemiological data (occurrence record) as well as molecular data (genetic sequences), to recover the dynamic of an outbreak.

In the context of macroevolution, the present work contributes to the ongoing effort towards bridging the gap between inferences made from the fossil record and inferences made from contemporary data. It can be seen as an extension of the birth-death process with sampling through time (Stadler, 2010), allowing one to take into account fossil occurrences which evolutionary relationships to other taxa are not well resolved. However, we need to point out here that we assume removal of a sampled individual from the population, while a species continues to exist if a specimen is preserved in and observed from the fossil record. Numerical rather than our analytic treatment of the model can deal with non-removal upon sampling (Vaughan et al., 2019).

Many extensions of this model are possible. One of the simplest would be to consider time-varying rate parameters *λ*_*t*_, *µ*_*t*_, *ψ*_*t*_, *ω*_*t*_. The most widely used approach to do so in the recent literature has been to consider so-called *skyline* versions of birth-death processes (Stadler et al., 2013), meaning that the rate parameters are piecewise constant functions. Because we expressed all our results based on two key functions *u*_*t*_ and *p*_*t*_, whose analytical expressions can easily be obtained considering piecewise constant rates, our central result would still hold. The challenge, however, would be to use such a model with an appropriate number of rate shifts so as not to overfit the data, although in a Bayesian context the number of rate shifts could also be estimated as part of a reversible jump MCMC scheme (Green, 1995).

Another important extension of the model would be to allow for the possibility not to remove individuals upon sampling. In fact, the model considered in Stadler (2010) does assume that *ψ*-sampled individuals keep living in the process. However, it is central for the derivation of our result that *ω*-sampled individuals are removed upon sampling. Our choice to remove all individuals upon sampling also seems reasonable for a number of applications in epidemiology, since sampled individuals are generally those that have sought medical attention, and -even if not quarantined - might be much less likely to spread the disease (but see also (Gavryushkina et al., 2014)).

Recently, Stadler et al. (2018) introduced an other extension of the birth-death model with sampling through time to account for data on the lifetime of some individuals. It assumes that individuals can be sampled multiple times, and that the first and last sampling event are recorded along the tree. This extension of the model could be transposed in our setting, where the record of occurrences would become a record of sampled lifetime intervals. Such an approach would allow to use infection interval data or stratigraphic range data.

In summary, we present a way to analytically calculate the likelihood of phylogenetic trees together with occurrence data. So far, this likelihood could only be calculated analytically for the phylogenetic tree or for the occurrence data. Thus, we view this work as a major step towards coherent statistical analysis of different data sources under the birth-death model with sampling through time.

## Acknowledgements

The authors are very grateful to Rachel Warnock for helpful comments on potential applications of the model.

### Appendix

#### Proof of Proposition 2.1

Note that reversing the direction of time in (2.1) we can write the ODE for *u*(*t*_*or*_ − *s*) as

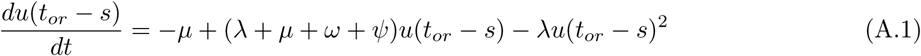

Applying the generator 𝔸_*k*_ on function *f*_*k*_(*s, i*) we obtain

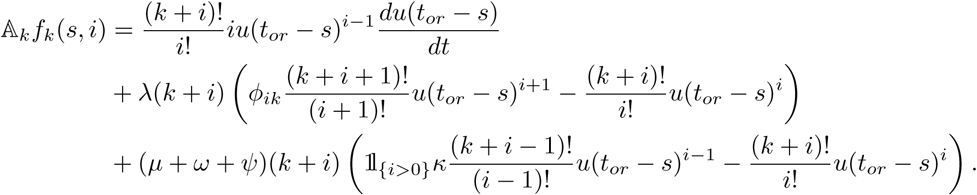

Since *i* ≥ 0, noting that

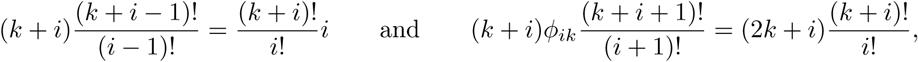

and using (A.1) we get

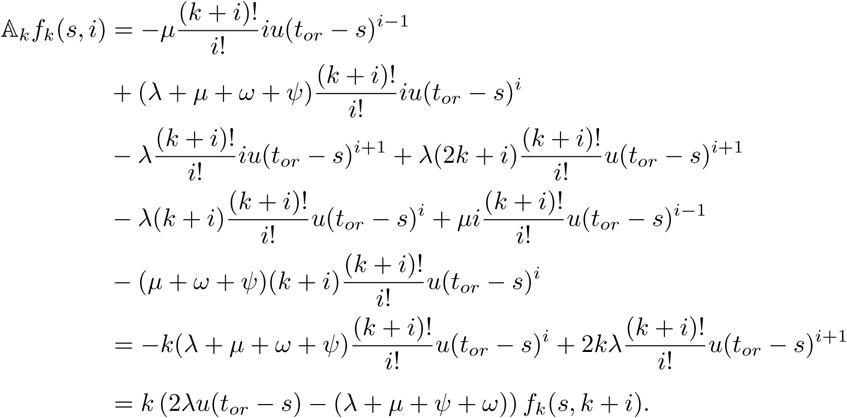

We now prove that *M*_*s*_ defined by (2.7) is a martingale w.r.t. the filtration ℱ_*s*_ generated by process *X*. Note that since this process has generator 𝔸_*k*_ the following is a ℱ_*s*_-martingale (see Chapter 4 in Ethier and Kurtz (1986))

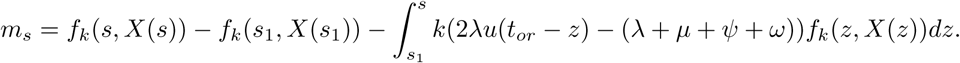

Writing this equation in differential form we obtain

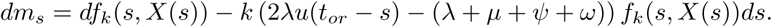

Multiplying both sides by the integrating factor

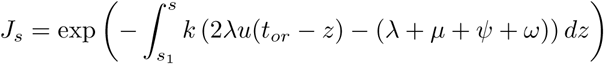

we get

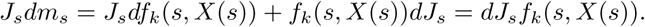

Upon integration we see that 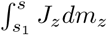 is a martingale which implies that *J f* (*s, X*(*s*)) is a positive ℱ_*s*_-martingale in the time-interval [*s*_1_, *s*_2_]. Note that *J*_*s*_ satisfies the ODE

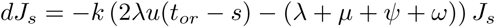

which is similar to (2.2). Exploiting this similarity allows us to write

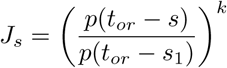

Now the fact that *J*_*s*_*f*_*k*_(*s, X*(*s*)) is a positive ℱ_*s*_-martingale proves that *M*_*s*_ is also such a martingale. This completes the proof of this proposition. □

#### Proof of Lemma 4.1

Observe that

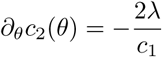

and all higher order derivatives of *c*_2_(*θ*) are zero, i.e. 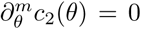 for all *m* ≥ 2. As *q*(*t, θ*) is a simple quadratic function of *c*_2_(*θ*) we get from chain-rule for derivatives that

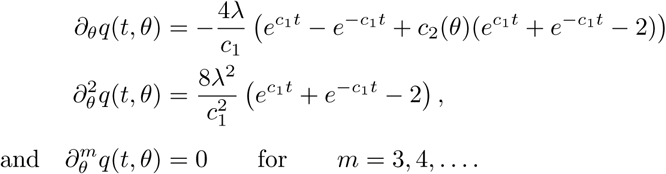

As *q*(0, *θ*) = 4 is independent of *θ* we can express the derivatives of *R*_*k*_(*t, θ*) as

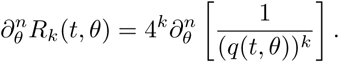

Using the derivatives of *q*(*t, θ*) computed above and the *Faà di Bruno’s* formula (see Fraenkel (1978)) we obtain the expression for 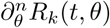 reported in part (A).

For part (B) observe that we can write *u*(*t, θ*) as

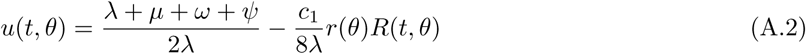

Where

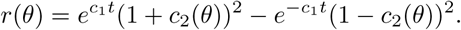

The derivatives of *r*(*θ*) can be obtained as

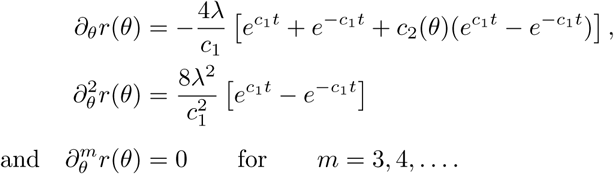

Applying the generalized product-rule for derivatives to formula (A.2) we can express 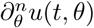 as

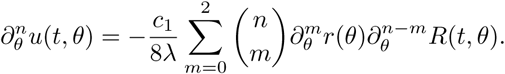

Substituting the values of 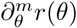 and simplifying, we obtain the expression for 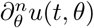 reported in part (B). This completes the proof of this lemma. □

